# Tracing the origins of molecular signals in food through integrative metabolomics and chemical databases

**DOI:** 10.1101/2025.10.21.683649

**Authors:** Alejandro Mendoza Cantu, Julia M. Gauglitz, Wout Bittremieux

## Abstract

Foods contain thousands of chemical constituents beyond macronutrients, including bioactive metabolites, processing by-products, and contaminants that remain poorly characterized. The Periodic Table of Food Initiative (PTFI) is establishing a standardized global reference for food composition using untargeted mass spectrometry. We analyzed the first PTFI release (∼24,000 molecular features across 500 foods) by linking annotated and unannotated signals to curated databases of pharmaceuticals, agrochemicals, food contact chemicals, and natural products. Annotated compounds revealed characteristic chemical patterns across food groups, while unannotated features exposed xenobiotic signatures and potential contamination pathways. A taxonomy-aware search identified unexpected natural products, such as biochanin A and phlorizin produced by Canada thistle. Together, these analyses show how agricultural practices, environmental exposures, and processing shape food chemistry and highlight the value of food metabolomics for advancing a One Health understanding of the molecular connections between the environment, food systems, and human health.

## Introduction

Food is one of the most direct and complex interfaces between humans and their environment. It provides essential nutrients, mediates chemical exposures, and reflects agricultural and processing practices. Understanding the molecular composition of food is therefore important not only for health and nutrition, but also for assessing environmental and industrial influences on the food supply.

While foods are typically described in terms of macronutrients—such as protein, fat, and carbohydrates—this represents only one view of their full chemical complexity. In reality, foods contain thousands of additional compounds, including bioactive molecules, processing by-products, and potential contaminants, many of which remain poorly characterized.^1^

Despite advances in analytical chemistry, comprehensive molecular profiles are still unavailable for most foods.^1^ This lack of information limits our ability to assess dietary exposures, evaluate food quality, and understand the broader health implications of food composition. Better molecular-level data could support more accurate nutritional recommendations, improved food safety monitoring, and the discovery of beneficial natural compounds. Existing food composition databases^2–4^ are often limited in scope, focusing on industrially processed or widely commercialized foods, with minimal representation of regionally important, traditional, or under-studied items.

Untargeted liquid chromatography mass spectrometry (LC-MS)-based metabolomics allows for broad detection of both known and unknown metabolites, making it especially valuable for exploring poorly characterized foods. Untargeted metabolomics initiatives like the Global FoodOmics Project^5^ and the Food Metabolome Repository^6^ yield insights into food by providing publicly available raw MS data, revealing the chemical complexity in food beyond compositional analysis.

To complement these ongoing efforts to understand what is in our food, the Periodic Table of Food Initiative (PTFI) was established.^7^ The PTFI is a global research effort to develop standardized protocols for untargeted metabolomics analysis of foods, with the goal of building a comprehensive, publicly accessible reference of food composition. It combines high-resolution LC-MS profiling with structured metadata based on ontologies like FoodOn,^8^ enabling scalable and interpretable foodomics analyses. As an initial release, the PTFI has made data for 500 commonly consumed foods available. These data include quantitative measurements for thousands of molecular features, covering both known reference compounds and a large set of unknown signals. As is typical in untargeted metabolomics, a substantial portion of the detected features remain unannotated, presenting an opportunity to explore their origins using external chemical databases.

In this study, we analyzed the PTFI dataset by matching both annotated and unannotated molecular features against curated databases of natural products, pharmaceuticals, agrochemicals, and food contact chemicals. This enabled us to examine the likely origins of chemical signals in food, identify potential contaminants, and detect unexpected sources of known natural products. By tracing these molecular linkages between biological, environmental, and industrial domains, our analysis contributes to a One Health understanding of how interconnected systems shape the chemical composition of the global food supply. These examples illustrate how knowledge bases can help interpret complex metabolomics signals.

## Results & discussion

The PTFI untargeted LC-MS dataset includes 500 globally relevant foods like grains, fruits, vegetables, meats, and dairy collected from various U.S. locations. Plant foods are the most represented group, spanning 70% of the data, followed by animal food products (15%), and the remaining percentage corresponding to algal and fungal products. The data consist of 24,721 unique metabolic features, which were annotated against PTFI in-house reference libraries and subsequently split into structurally annotated and non-structurally annotated features for analysis, allowing us to apply a two-pronged analysis strategy (**Figure 1a**). First, we cross-referenced the set of structure-annotated compounds with curated knowledge bases to infer likely functional classes and potential exogenous sources (**Figure 1b**). Of the 900 structure-annotated compounds, a subset matched external databases of drugs, agrochemicals, and food contact chemicals. Furthermore, we performed a taxonomy-aware screen to identify putative novel producers of known natural products.

**Figure 1.**
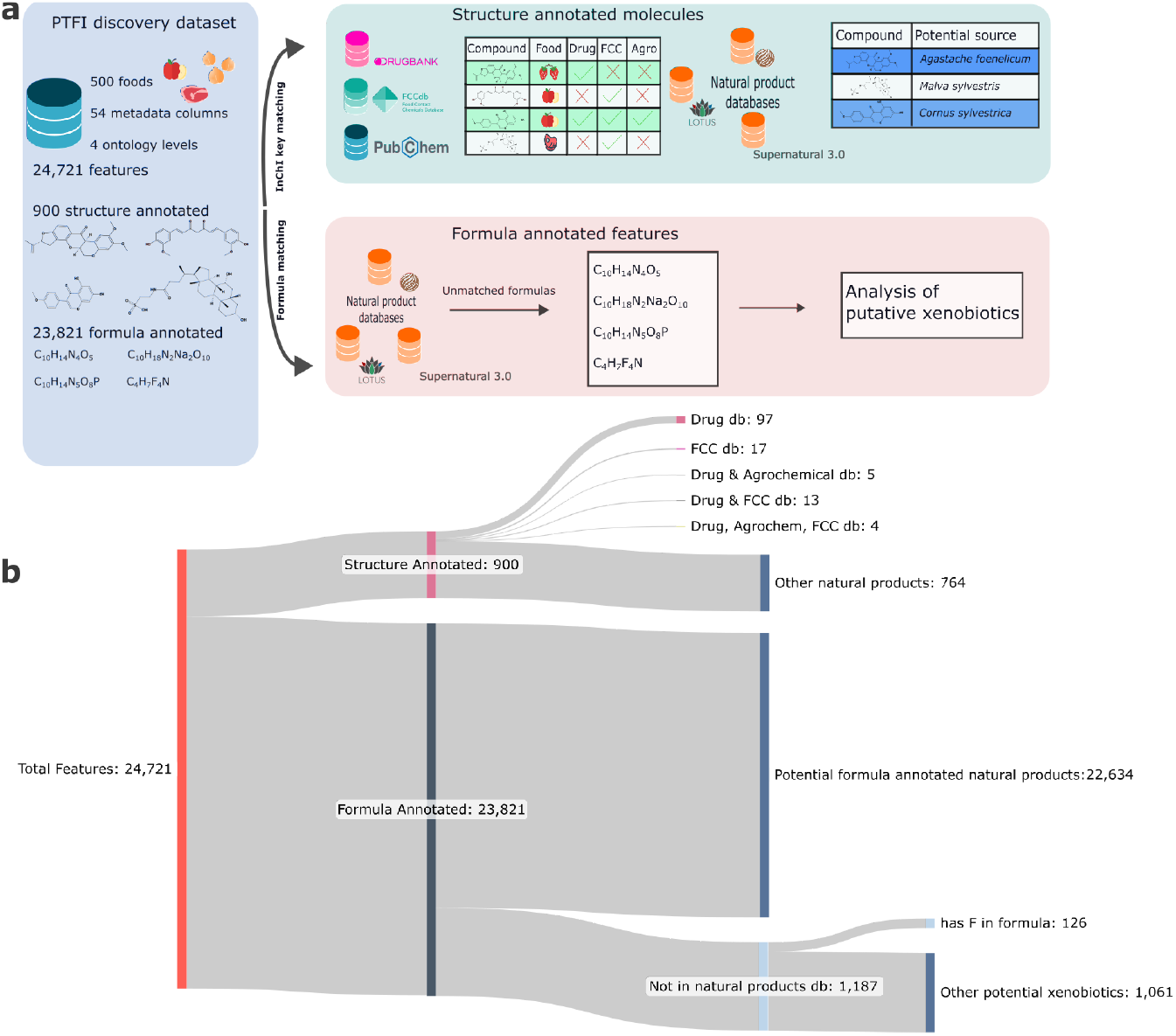
Analysis workflow and categorization of molecular features in the PTFI dataset. **(a)** Schematic of the data analysis workflow. Structure-annotated molecules were matched by InChI key to curated databases of pharmaceuticals, agrochemicals, and food contact chemicals to infer potential sources. Formula-annotated features were screened by molecular formula against natural product databases to identify formulas not found in those references; these were classified as putative xenobiotics. **(b)** Sankey diagram illustrating the categorization of 24,721 molecular features detected across 500 foods, subdivided by chemical characteristic or source of annotation.

Second, for non-structurally annotated features, molecular formulas were provided to enable downstream matching. We used formula-level filtering to prioritize unannotated features that are unlikely to be biogenic and may represent xenobiotics (**Figure 1b**). Of the 23,821 features with only formula annotations, 22,634 match a natural product molecular formula, indicating that they may have potential pharmacological, agrochemical, or contaminant origins. In addition, 1,187 features were absent from the natural product reference databases and flagged as potential xenobiotics. This xenobiotic subset was further prioritized by screening for formulas containing fluorine.

### External database matching identifies contaminants, bioactives, and therapeutic agents in foods

To obtain a high-level view of chemical classes with potential regulatory or toxicological relevance, we summed signal intensities for compounds matched to DrugBank, FCCdb, and the PubChem subset within each FoodOn level 1 category (**Figure 2a**). Plant food products showed the highest aggregate signal for DrugBank-matched compounds, consistent with the breadth of phytochemicals that have direct pharmacological activity or overlap structurally with the pharmacologically relevant space.^9^ Animal food products exhibited the next-highest DrugBank signal, which may reflect endogenous bioactive metabolites, feed-derived exposures, veterinary drugs carryover, or processing-related carryover.

**Figure 2.**
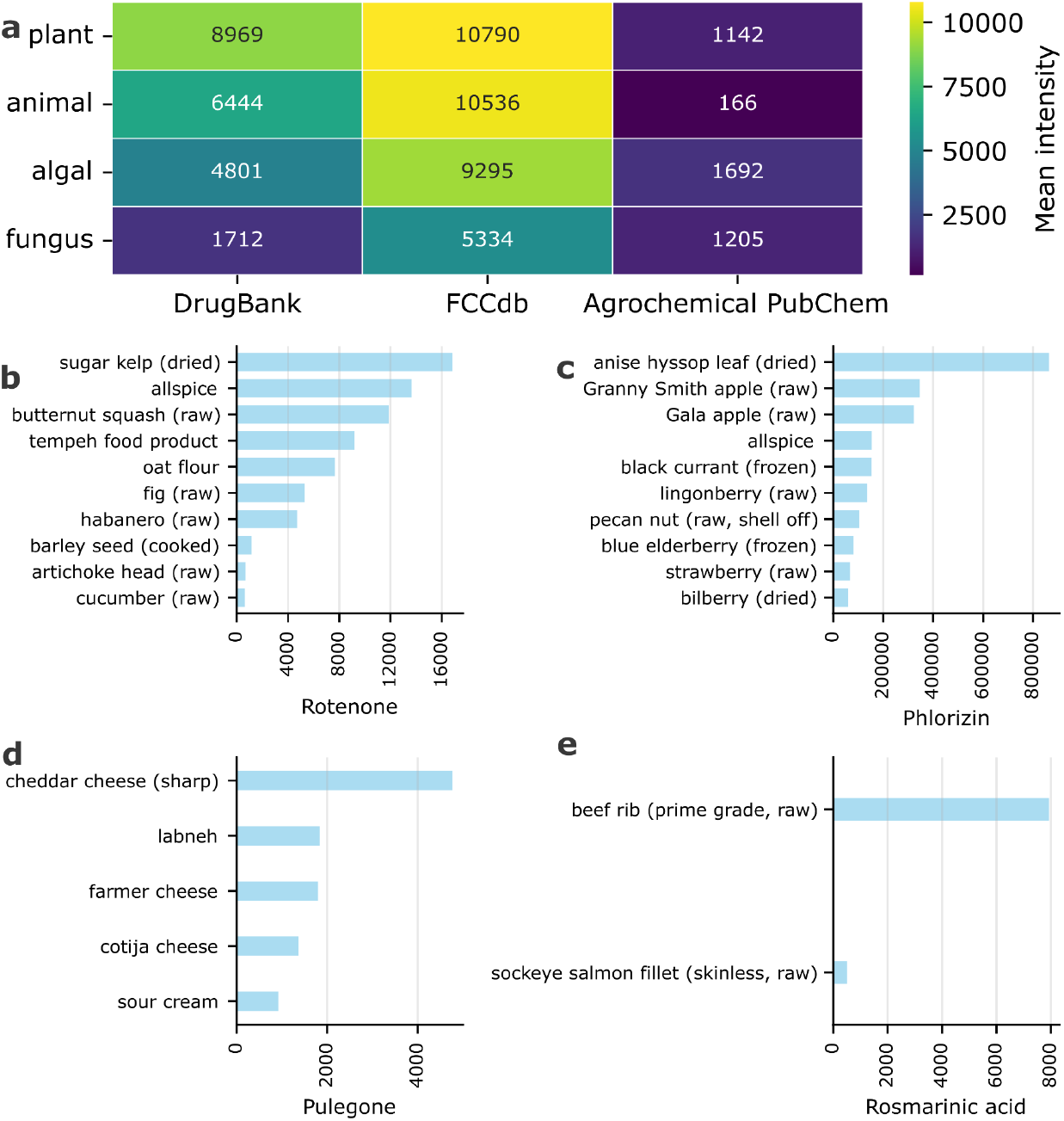
Analysis of annotated metabolites matched to external drug, contact chemical, and agrochemical knowledge bases. **(a)** Heatmap showing the mean intensity of features from the four principal PTFI ontologies (plant, animal, algal, fungus) across three databases: DrugBank, FCCDdb, and Agrochemical PubChem. **(b–d)** Bar plots showing the signal intensity of selected compounds matched to external databases across their up to top 10 food sources for **(b)** the piscicide rotenone, **(c)** the bioactive compound phlorizin, and **(d)** the contact chemical pulegone. **(e)** Bar plot showing the signal intensity of rosmarinic acid in animal products.

The FCCdb-associated molecular signal was consistently high across animal, plant, and algal food products, in contrast to the much lower agrochemical matches observed across all categories. This pattern reflects both the fundamental difference in what these databases capture and their relative sizes: FCCdb contains 35 matched compounds in our dataset compared to only nine agrochemical matches. The top FCCdb matches include linoleic acid in plant products, hexadecanoic acid in animal products, and azelaic acid across multiple food categories; all naturally occurring, abundant fatty acids which are common components of food matrices.^10–12^ While FCCdb was designed to catalog food contact chemicals and additives, it does not distinguish between natural food constituents and industrially produced chemicals.

In contrast, agrochemical matches were dominated by naturally occurring compounds with pesticidal properties (piperine, rotenone), agrochemical precursors (4-hydroxybenzoic acid), and plant growth regulators (indoleacetic acid, folic acid). The agrochemical signal was lowest in animal products, which is consistent with the expectation that most agrochemical compounds originate from plant sources due to agricultural activity or environmental contamination, rather than being intrinsic to animal food matrices.

Next, we highlight four compounds which matched to the selected chemical databases to illustrate how database-informed comparison reveals diverse possible routes of chemical entry into foods. We selected rotenone for its prominence among agrochemical matches and known health implications, phlorizin for its unexpected distribution beyond documented sources, pulegone for its dual nature as both a natural product and regulated contaminant, and rosmarinic acid as an example of plant-derived bioactives in animal products. From these cases we hypothesize intentional application, environmental contamination, and processing-related transfer.

Rotenone, a naturally occurring isoflavonoid found in non-edible parts of some plant species, has insecticidal and piscicidal applications,^13^ and was among the most prominent agrochemical matches in the dataset. The top 10 food sources for this compound included both aquatic and terrestrial products (**Figure 2b**), indicating its broad presence. Sugar kelp showed the highest mean normalized abundance, which aligns with prior exploration of rotenone as an algal crop protective agent.^13^ In terrestrial contexts, rotenone was prevalent in vegetables and herbs/spices, consistent with its continued use on certain crops despite regulatory restrictions in some regions.^14^ Particularly notable was its detection in organic oat flour; an unexpected finding, as rotenone is a prohibited substance under U.S. National Organic Program standards.^15^ Possible explanations include environmental drift from nearby conventional farms, persistence of residues in soil from prior applications, or non-compliance. The documented association of rotenone exposure with Parkinson’s disease^16^ underscores the relevance of such observations, once validated, for food safety monitoring.

Phlorizin is a dihydrochalcone primarily reported in apples and other members of the *Malus* genus with reported antidiabetic, antioxidant, and anti-inflammatory activities.^17^ While two kinds of apples are among the top foods where phlorizin was detected, the highest concentrations were actually found in anise hyssop (**Figure 2c**), possibly playing an unnamed role in its use in traditional medicine.^18^ Additionally, phlorizin was also detected in several other non-apple plant products such as allspice, pecan nut, and a number of different berries. This finding suggests either a broader biosynthetic distribution than currently described in the literature or transfer during multi-ingredient processing.

Pulegone is a cyclic monoterpenic compound primarily found in plants of the Lamiaceae family, such as mint (*Mentha* spp.) and thyme (*Thymus* spp.).^19^ Some regulatory bodies have defined it as a hazardous substance; for example, it is listed as a carcinogen under California’s Proposition 65. While the PTFI dataset does not include mint and thyme, it contains several other Lamiaceae herbs (basil, lemon balm, anise hyssop, and beautyberry). However, our analysis revealed that pulegone was detected exclusively in dairy-based products, specifically three types of cheese, labneh, and sour cream (**Figure 2d**). Two plausible sources could explain this observation. First, pulegone’s antimicrobial properties make it an occasional component in cheese coatings designed to extend shelf life, from which it could migrate into the food.^20,21^ A second possibility is carryover from animal feed, as it has been reported that aromatic compounds from essential oils in cattle diets, derived from pulegone-producing plants, can be transferred to milk and concentrated in cheese.^22^ This latter pathway would also explain its presence in products not typically coated, such as labneh and sour cream.

As a final case study motivated by the search for bioactive compounds in animal sources, we present the finding of rosmarinic acid (**Figure 2e**). This compound was only found in two animal products in the dataset: beef rib and sockeye salmon. This is a bioactive compound known for its beneficial effects in treating ailments like cancer and diabetes.^23^ Its detection in fish and beef meat might be explained by its intentional addition as a natural antioxidant, as it is a common additive used to prevent lipid oxidation in meat products.^24^ Furthermore, these are not just spurious amounts introduced as trace environmental contaminants. The levels of rosmarinic acid detected in beef (signal intensity 7,932) are higher than in some plants, such as purslane leaf (signal intensity 5,890), although there are also exceptionally high abundances observed in other plants, such as holy basil (signal intensity >5,000,000).

Together, these case studies highlight two complementary strengths of our analysis: the ability to flag potential contamination events that merit regulatory or analytical follow-up, and the capacity to uncover unexplored sources of bioactive molecules.

### Presence of DrugBank features across food types

Building on the previous examples of agrochemical contamination, unexpected bioactive distributions, and food contact chemical transfer, we extended our analysis to systematically examine DrugBank therapeutic categories across food types. We selected 20 groups of interest from DrugBank’s therapeutic categories, which classify drugs based on their use in treatment and therapy. The analysis revealed plant foods dominating all categories, as expected given their phytochemical diversity (**Figure 3a**). However, certain categories showed unexpected animal product signals. Anticarcinogenic agents, anticoagulants, and phytoestrogens all appeared in animal foods despite not being endemic to these matrices, pointing to potential carry-over via the food chain. Cross-referencing the annotated molecules with DrugBank therapeutic categories enabled identification of potential drug residues and bioactive compound sources in the food supply. To illustrate these transfer mechanisms, we examined three compounds that exemplify feed-to-animal pathways: 4-hydroxycoumarin (anticoagulant), daidzein (phytoestrogen), and salicylic acid (anti-inflammatory agent).

**Figure 3.**
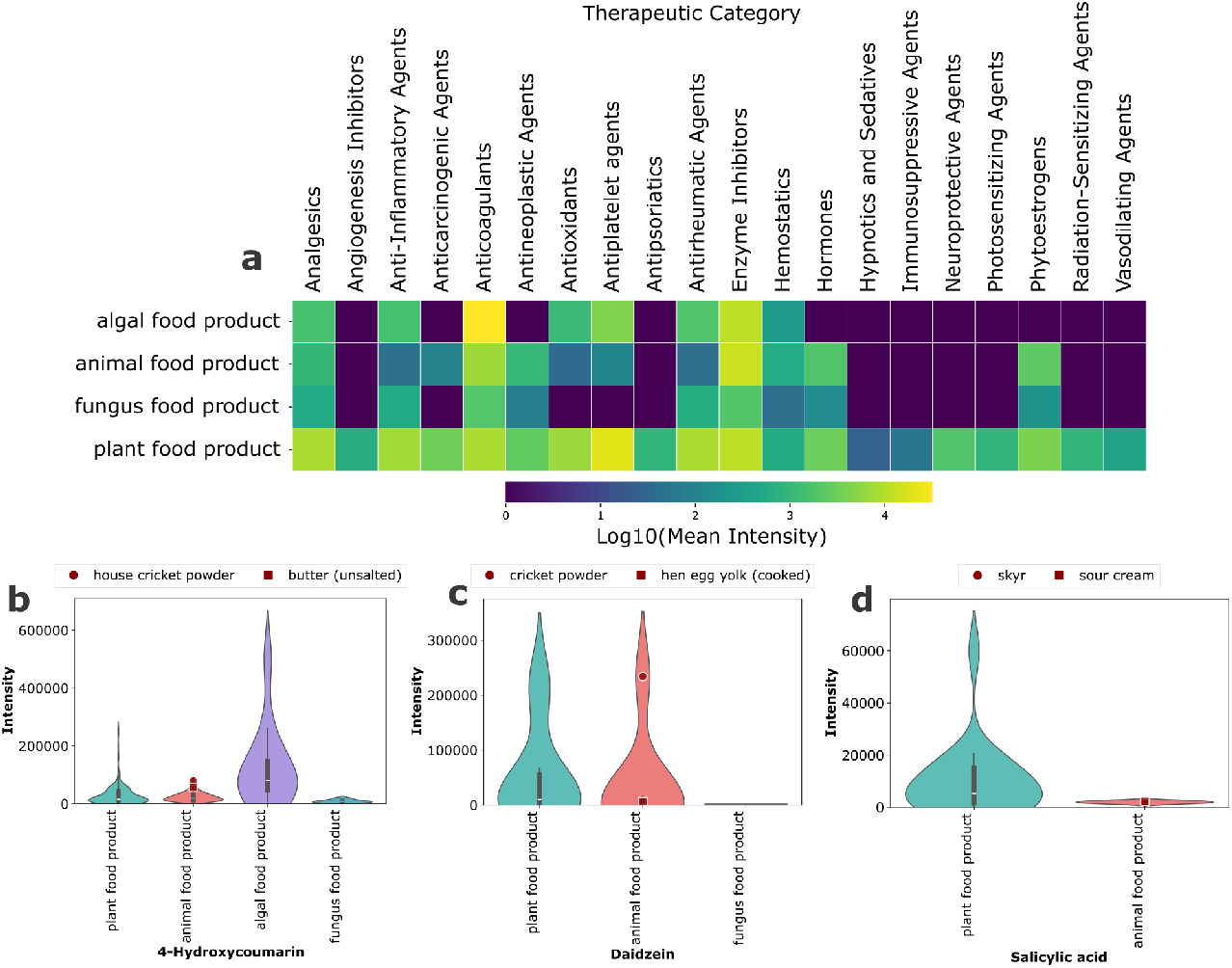
Distribution of DrugBank therapeutic categories and veterinary-approved drugs across foods. **(a)** Heatmap showing the log-transformed mean intensity of compounds grouped by therapeutic category across FoodOn level 1 food groups. **(b–d)** Violin plot showing the abundance of **(b)** 4-hydroxycoumarin, **(c)** daidzein, and **(d)** salicylic acid across food categories.

4-hydroxycoumarin, a microbially produced anticoagulant^25^ as well as found as a product of fermentation and in some plant biosynthetic pathways,^26,27^ was most commonly detected in algal food products (**Figure 3b**). The elevated coumarin presence may be attributed to their photoprotective role, as coumarins have been shown to serve as UV-protective compounds in macroalgae, accumulating in cell walls and vacuolar membranes to absorb harmful radiation.^28^ Furthermore, it appears in 97% of animal food samples (63 of 65) and 97% of plant samples (194 of 200), with comparable intensities across both food categories. Two animal products which show the highest intensity are butter and (house) cricket powder, demonstrating its presence across diverse animal-derived matrices. The presence of 4-hydroxycoumarin in animal products is notable because it is not approved as a veterinary drug, ruling out pharmaceutical administration. This nearly ubiquitous presence at similar abundance levels in both animal and plant products suggests transfer from plant-based feeds or other dietary sources, or possibly carryover during mass spectral data collection.

Daidzein, classified under DrugBank’s phytoestrogen category, showed accumulation in animal products despite its primary association with soy-based plant sources. While present in hen eggs, it is substantially accumulated in (house) cricket powder (**Figure 3c**), possibly due to the fact that insects are known to efficiently absorb and metabolize dietary phenolic compounds from plant-based feeds.^29^ Furthermore, daidzein has also been used as a supplement to improve egg quality, as demonstrated in laying hens where dietary daidzein supplementation (30 mg/kg) showed reduced cholesterol deposition in egg yolk through modulation of lipid metabolism.^30^

Salicylic acid, which matched DrugBank’s anti-inflammatory and analgesic categories, was detected in dairy products including skyr and sour cream (**Figure 3d**). While this compound is a known anti-inflammatory agent, its presence in food is noteworthy for individuals with salicylate intolerance. The detected residues (skyr: signal intensity 1,533; sour cream: signal intensity 2,295) are comparatively low, representing only 2.5–5.0% of the levels found in salicylate-rich plant sources such as lemon balm leaf (signal intensity 60,000) or hemp seed (signal intensity 18,600). While one potential source is its natural occurrence in animal feed like alfalfa, the transfer of salicylic acid from feed to milk in ruminants is considered very limited and unlikely.^31^ Given the relatively low concentrations, the origin of these residues remains uncertain and could include multiple pathways: natural occurrence in feed with minimal transfer, veterinary treatments such as aspirin administration, or biocide residues from processing or transfer of salicylates from fermented silage, as it has been found that salicylates may be present in silage fed to dairy cattle.^32^

Importantly, as this analysis is limited to compounds annotated through the PTFI reference library of authentic standards, the detection of pharmacologically relevant compounds in animal products may represent only a subset of signatures from carry-over compounds and transfer mechanisms.

### Discovery of underreported bioactive natural product producers

To identify unexpected biological sources of bioactive natural products, we compared the genus of each compound in the PTFI dataset that matched pharmaceutical databases (DrugBank or PubChem Drugs) against genus-level occurrence records in the aggregated LOTUS, COCONUT, and SuperNatural databases. This procedure yielded 776 compound–genus pairs for which the producing genus was absent from all reference sources. To focus on specialized metabolites rather than ubiquitous compounds, we excluded any metabolite detected in more than 10% of all samples, retaining 84 novel producer candidates (plant genus) and 243 compound–genus pairs.

To contextualize the observation of known compounds in novel food matrices, each compound was assigned a chemical class using ClassyFire,^33^ with the most common sample name used for identification (**Figure 4a**). The results highlight several species as particularly rich in unreported compounds, such as carrot (*Daucus carota*) and soybean (*Glycine max*), with 30 and 20 compounds, respectively. While soybean shows a high total count of unreported molecules, the majority of these molecules are in one class, prenol lipids. In contrast, carrot (*Daucus carota*), Canada thistle (*Cirsium arvense*), purslane (*Portulaca oleracea*), and common mallow leaf (*Malvia sylvestris*) present a more diverse novel compound content, notably containing a significant proportion of flavonoids that are not commonly reported for these foods (**Figure 4a**).

**Figure 4.**
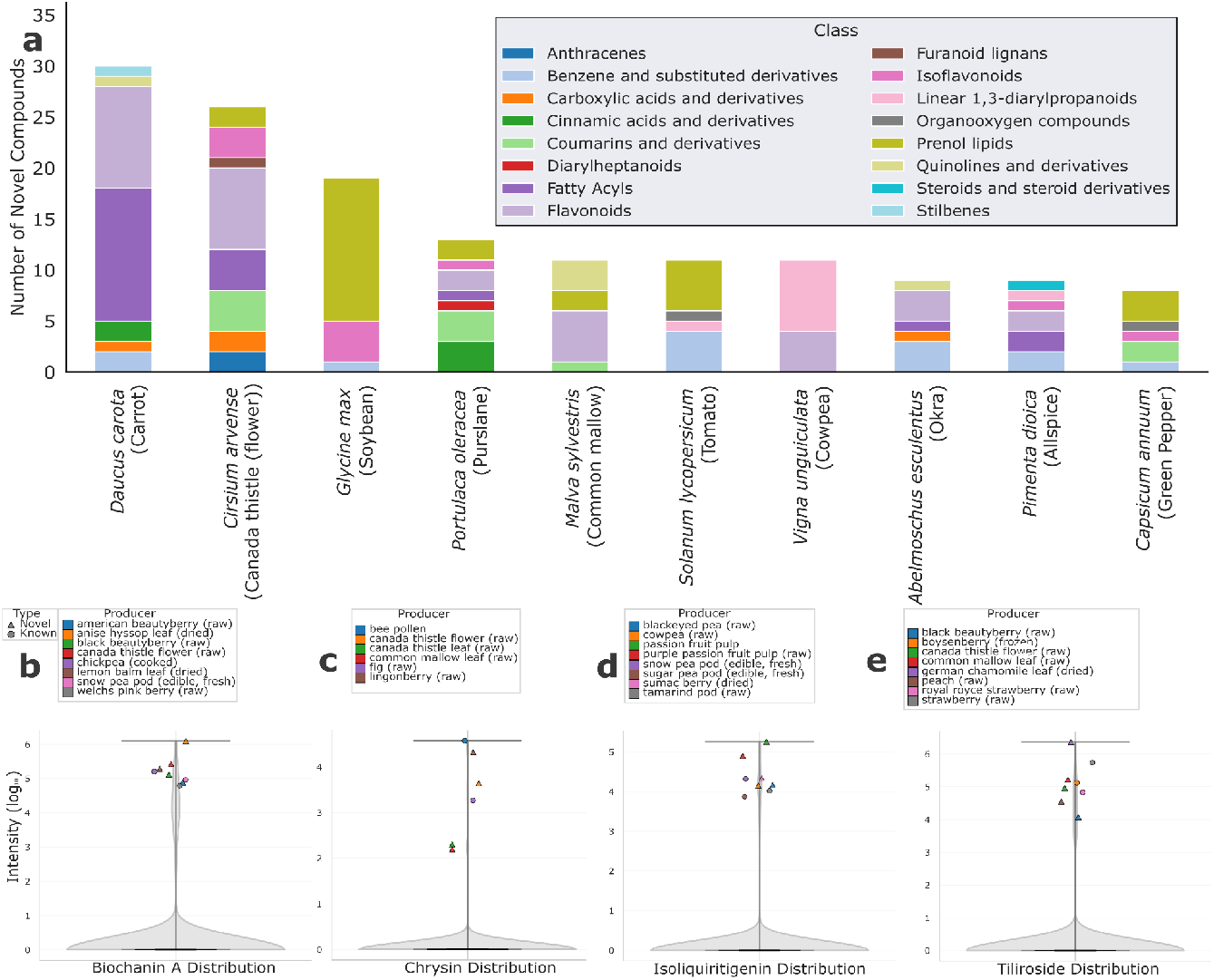
Analysis of novel plant natural product producers. **(a)** Top 10 species by number of unique unreported metabolites discovered, categorized according to chemical classes determined by ClassyFire. **(b–e)** Top putative novel sources for **(b)** biochanin A, **(c)** chrysin, **(d)** isoliquiritigenin, and **(e)** tiliroside, showcasing the intensity against the most abundant known producers of these compounds.

After mapping the overall diversity of the unreported screened compounds, we focused on specific examples to highlight our findings. The compounds discussed below (**Figure 4b–e**) were selected both for their known bioactivity and therapeutic properties,^34–37^ and because they represent discoveries where a novel producer showed a remarkably high abundance, often surpassing all known sources in the database.

For instance, biochanin A is an isoflavone normally found in chickpeas with diverse therapeutic applications, such as neuroprotective and anticancer properties.^34^ Notably, anise hyssop (*Agastache foeniculum*) was not only identified as a novel source but also emerged as the top producer of this metabolite, with an intensity far exceeding all other foods analyzed (**Figure 4b**).

Similarly, we observed a compelling pattern for isoliquiritigenin (**Figure 4c**), a chalcone with antitumoral and antioxidant activities.^35^ The two highest concentrations were found in the pulp of two passion fruit (*Passiflora edulis*) variants. This consistency across samples from the same botanical family strengthens the evidence that these are genuine biosynthetic findings rather than experimental artifacts.

A similar trend was observed for chrysin, a flavonoid typically associated with honey and bee pollen, where hepatoprotective and antidiabetic properties have been reported.^36^ In the PTFI data, the highest intensities were detected in two parts of the same plant, the flower and leaf of Canada thistle (*Cirsium arvense*), with the concentration in the flower being considerably higher than in the leaf (**Figure 4d**).

Finally, a comparable trend was observed for tiliroside, a dietary flavonoid with documented antiproliferative and antimicrobial activities.^37^ While tiliroside is commonly associated with strawberries, Canada thistle (*Cirsium arvense*) emerged as the top producer in our dataset, with additional detections in peach (*Prunus persica*) and common mallow leaf *(Malva sylvestris*) (**Figure 4e**). The consistency of Canada thistle as a high producer across multiple bioactive compounds and high count of unreported discoveries suggests that this species may represent an underexplored source of therapeutic natural products.

### Molecular formula annotated features reveal putative xenobiotics

The unannotated portion of the dataset comprised ∼24,000 molecular features for which only elemental formulas were available. We screened these formulas against the combined reference space of LOTUS, COCONUT, and SuperNatural to identify candidates unlikely to originate from known natural products (**Figure 1**). A total of 1187 features (4.8%) were absent from the natural product formula set and were therefore classified as putative xenobiotics. To further enrich for likely anthropogenic compounds, we applied an elemental filter targeting formulas containing fluorine. Fluorine-containing natural products are exceptionally rare and thus strong candidates for anthropogenic xenobiotics in food matrices, as they are prevalent in synthetic agrochemicals, industrial additives, and persistent pollutants.^38^

The distribution of fluorine-containing xenobiotics across FoodOn level 1 categories revealed distinct xenobiotic fingerprints for major food groups (**Figure 5a**). Within the animal category, subcategory-level clustering uncovered profiles for specific product types (**Figure 5b**). For example, there is a distinct grouping of dairy products characterized by a unique combination of fluorinated formulas. At the level of molecular formula annotation it is not possible to group these features into chemical categories, though several potential sources likely contribute to these signals. The fluorinated signals observed could originate from veterinary pharmaceuticals and pesticide residues used in dairy production. Additionally, a class of fluorinated compounds that have reported health consequences and which are now ubiquitous in our environment are per- and polyfluoroalkyl substances (PFAS), which may be a contributing factor to these detections. While there have been reports of PFAS contamination in dairy products,^39^ the distinct fluorine patterns in these dairy products indicate that comprehensive screening may be necessary to fully assess xenobiotic exposure from dairy consumption.

**Figure 5.**
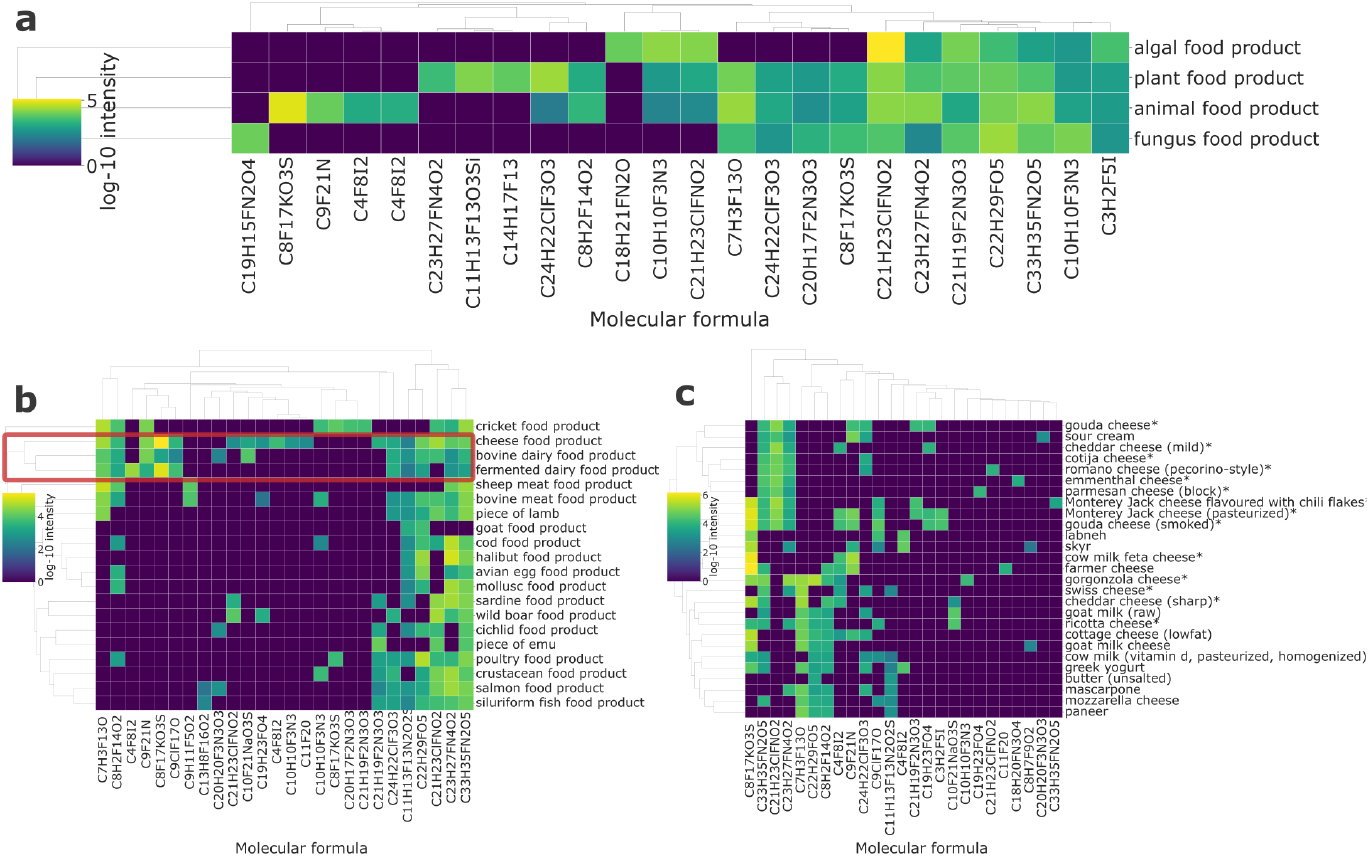
Fluorine-containing features reveal the presence of potential xenobiotic unknowns. **(a)** Clustered heatmap of top 10 fluorine-containing putative xenobiotics for each FoodOn level 1 category. **(b)** Clustered heatmap of top 10 fluorine-containing putative xenobiotics for each FoodOn level 2 animal product categories. A distinct group of dairy foods is indicated by the red box. **(c)** Clustered heatmap of top 10 fluorine-containing putative xenobiotics within dairy products. Hard cheeses are marked with an asterisk (*).

A deeper analysis demonstrated that even within this specific set of dairy samples, xenobiotic profiles can be highly differentiated (**Figure 5c**). Hard, aged cheeses share a core set of three recurring fluorinated features (C_33_H_35_FN_2_O_5_, C_21_H_23_ClFNO_2_, C_23_H_27_FN_4_O_2_) that were absent or low-abundance in fresh cheeses, suggesting that aging processes, protective coatings, or packaging materials used for hard cheeses may introduce specific fluorinated contaminants not found in fresh dairy products.

One prominent feature in the dairy cluster had the formula C_8_F_17_KO_3_S, potentially corresponding to the potassium salt of perfluorooctanesulfonic acid (PFOS), a well-documented PFAS contaminant in milk.^39^ This feature was consistently detected in milk but appeared at higher intensities in aged cheeses. The increase may be a concentration effect driven by the higher protein and fat content of cheese; the amphiphilic nature of PFOS causes it to bind to fats and proteins,^40^ which become significantly more concentrated as water is removed during maturation. Inspection of the compositional PTFI dataset, which accompanies the metabolomics data, shows that a cheese can greatly increase the protein content from 3.37g/100g in milk to over 22.90g/100g in Gouda.^41,42^

In conclusion, by matching the first release of the PTFI’s food metabolomics data with multiple external chemical and natural product knowledge bases, we sought to understand the origin of molecular signals in food data. Our complementary analyses highlight the interplay of agricultural practices, environmental exposures, biological biosynthesis, and industrial processing in shaping the molecular composition of foods. This systems-level perspective aligns with the One Health concept, which emphasizes the interdependence of human, animal, and environmental health. By tracing chemical linkages across these domains, food metabolomics can contribute to identifying routes of exposure, monitoring contaminants, and uncovering beneficial compounds that connect agricultural and public health outcomes. Together, these findings demonstrate how curated chemical occurrence information can be used to identify contamination pathways and discover new food sources of bioactive compounds with potential nutritional or therapeutic value, supporting One Health-oriented efforts to build safer and more sustainable food systems.

While our results illustrate the value of combining standardized food metabolomics with curated external resources, further interpretation is constrained by the absence of tandem MS fragmentation data, which limits the confidence of structural annotations and the ability to resolve isomeric compounds. In addition to the provided metadata, more precise contextual data such as processing steps, packaging materials, and storage histories would further aid in discriminating between endogenous biosynthesis, environmental uptake, and contamination.

Despite these limitations, the approach outlined here is broadly extensible. As the PTFI dataset grows in scope and incorporates richer chemical and contextual information, the same computational approach can be scaled to monitor emerging contaminants, identify novel bioactive compounds, and build an increasingly complete molecular map of the global food system—supporting efforts in nutrition, food safety, biodiversity conservation, and natural product discovery.

## Methods

### Data and preprocessing

We analyzed the first public release of the PTFI dataset, which includes untargeted metabolomics measurements for 500 commonly consumed food items.^7^ Each sample is accompanied by structured metadata describing its source organism, production method, and food classification, the latter organized hierarchically using the FoodOn ontology.^8^

The PTFI team generated these data using high-resolution LC-MS with time-of-flight mass spectrometry, resulting in 24,721 metabolic features defined by their mass-to-charge ratio (*m*/*z*) and chromatographic retention time index. 900 features were annotated through accurate mass and retention time matching against a reference library of over 1,500 authentic standards analyzed with the PTFI experimental protocols.^43^ Additionally, molecular formulas were derived for “known unknowns” of consistently observed features (in at least two unique experiments of samples in the same ontological family) that remained unannotated. All intensity values in the PTFI dataset were normalized within each injection to the median intensity of 33 internal standards, enabling tracking of compound abundance across different samples and analytical runs. No MS/MS data are available.

All data analysis was conducted within the PTFI online runtime environment hosted by the American Heart Association’s Precision Medicine Platform. The data were originally provided as a wide-format abundance matrix with features as rows and samples as columns. We converted this into a long-format table, where each row corresponds to a metabolite–sample measurement.

### Database integration and matching strategy

To interpret the chemical origin of observed features, we cross-referenced annotated compounds with DrugBank (v6.0)^44^ for pharmaceuticals, the Food Contact Chemicals Database (FCCdb; v5.0)^45^ for packaging-related substances, and a filtered subset of PubChem (v2025)^46^ entries corresponding to agrochemicals (e.g. pesticides and fungicides, based on the “Agrochemicals” table of contents category) and pharmaceuticals (based on Medical Subject Headings (MeSH) classifications). 93 compounds matched to DrugBank, corresponding to 400 unique therapeutic category terms. To focus on biologically relevant functions, we excluded non-therapeutic classifications such as chemical descriptors and industrial uses. From the remaining terms, we selected 20 broad therapeutic categories (e.g. Enzyme Inhibitors, Antioxidants) that capture diverse pharmacological activities. To assess potential biological origins of known compounds, we further integrated three natural product databases: LOTUS (NPOC2021, February 2021),^47^ COCONUT (v2.0),^48^ and SuperNatural (v3.0).^49^ Matching was performed using InChI keys for annotated compounds and molecular formulas for unannotated features.

### Compound origin analyses

To identify potential novel sources of known natural products, compounds were matched to food samples based on genus-level taxonomy. The genus was extracted from each sample’s organism name metadata field and compared against the genus-level organism annotations in the three natural product databases (LOTUS, COCONUT, SuperNatural). A compound was flagged as a potential novel discovery if the plant genus of the food sample was not among the known producers listed in any of the external databases. To focus on rare or specialized metabolites, compounds detected in more than 10% of all plant samples were excluded. The final output was a list of compound–genus pairs not previously reported in available natural product literature.

We used ClassyFire (version 1.0),^33^ a hierarchical chemical taxonomy system that assigns compounds to categories based on their structural features, to classify compounds for the novel producer analysis. Each compound was classified using its InChI.

For features that remained unannotated, i.e. those for which only the molecular formula was known, a formula-based screen was used to identify putative xenobiotics. Molecular formulas for 23,821 such features were compared to the union of formulas from the three natural product databases (LOTUS, COCONUT, SuperNatural). Any formula not found in this reference set was classified as a putative xenobiotic. To further prioritize likely contaminants, we filtered for formulas containing fluorine (F), as this element is uncommon in biosynthetic products but frequently observed in industrial and agricultural compounds. The resulting list provided a set of candidate xenobiotic features for further analysis and interpretation.

## Data availability

The PTFI dataset analyzed in this study is available through the American Heart Association’s Precision Medicine Platform at https://pmp.heart.org/. External compound databases were retrieved from their corresponding web resources: LOTUS (vNPOC2021; https://lotus.naturalproducts.net/), COCONUT (v2.0; https://coconut.naturalproducts.net/), SuperNatural (v3.0; http://bioinf-applied.charite.de/supernatural_3/), DrugBank (v6.0; https://www.drugbank.ca/), Food Contact Chemicals Database (v5.0; https://foodpackagingforum.org/resources/databases/fccdb) and PubChem (v2025; https://pubchem.ncbi.nlm.nih.gov/).

## Code availability

All analyses were conducted in Python 3.12 within the online PTFI data analysis platform (https://pmp.heart.org/). Data processing was performed using the Pandas (v2.2.3)^50,51^ and Scikit-Learn (v1.7.0)^52^ libraries. Figures were generated using Matplotlib (v3.9.3)^53^ and Seaborn (v0.13.2).^54^ Analysis code, including notebooks for data processing, database matching, and figure generation, is available as open source at https://github.com/AlejandroMC28/ptfi_externaldb.

## Acknowledgements

This work was conducted using data provided through the PTFI 2024 Data Challenge. We thank the PTFI team and the American Heart Association’s Precision Medicine Platform for enabling access to the dataset and analysis environment. This work was supported by the Research Foundation – Flanders (FWO G0AGQ24N).

## Author contributions

A.M.C.: conceptualization, methodology, software, investigation, data curation, writing - original draft, writing - review & editing, visualization. J.M.G.: conceptualization, methodology, writing - original draft, writing - review & editing, supervision, funding acquisition. W.B.: conceptualization, methodology, writing - original draft, writing - review & editing, supervision, funding acquisition.

## References

1. Barabási, A.-L., Menichetti, G. & Loscalzo, J. The unmapped chemical complexity of our diet. Nat. Food 1, 33–37 (2019).

2. FooDB. https://foodb.ca/.

3. USDA FoodData Central. https://fdc.nal.usda.gov/.

4. Food composition data | EFSA. https://www.efsa.europa.eu/en/data-report/food-composition-data (2021).

5. Gauglitz, J. M. et al. Untargeted mass spectrometry-based metabolomics approach unveils molecular changes in raw and processed foods and beverages. Food Chem. 302, 125290 (2020).

6. Sakurai, N. et al. The Thing Metabolome Repository family (XMRs): comparable untargeted metabolome databases for analyzing sample-specific unknown metabolites. Nucleic Acids Res. 51, D660–D677 (2023).

7. Jarvis, A. et al. Periodic Table of Food Initiative for generating biomolecular knowledge of edible biodiversity. Nat. Food 5, 189–193 (2024).

8. Dooley, D. M. et al. FoodOn: a harmonized food ontology to increase global food traceability, quality control and data integration. Npj Sci. Food 2, 23 (2018).

9. Chihomvu, P., Ganesan, A., Gibbons, S., Woollard, K. & Hayes, M. A. Phytochemicals in Drug Discovery—A Confluence of Tradition and Innovation. Int. J. Mol. Sci. 25, 8792 (2024).

10. Showing Compound Azelaic acid (FDB012192) - FooDB. https://foodb.ca/compounds/FDB012192.

11. Showing Compound alpha-Linoleic acid (FDB006287) - FooDB. https://foodb.ca/compounds/FDB006287.

12. Showing Compound Hexadecanoic acid (FDB011679) - FooDB. https://foodb.ca/compounds/FDB011679.

13. El-Sayed, W. M. M. et al. Environmental influence on rotenone performance as an algal crop protective agent to prevent pond crashes for biofuel production. Algal Res. 33, 277–283 (2018).

14. Gupta, R. C. & Milatovic, D. Insecticides. in Biomarkers in Toxicology 389–407 (Elsevier, 2014). doi:10.1016/B978-0-12-404630-6.00023-3.

15. Federal Register :: National Organic Program; Amendments to the National List of Allowed and Prohibited Substances (Crops, Livestock and Handling).

16. Van Laar, A. D. et al. Transient exposure to rotenone causes degeneration and progressive parkinsonian motor deficits, neuroinflammation, and synucleinopathy. Npj Park. Dis. 9, 121 (2023).

17. Ehrenkranz, J. R. L., Lewis, N. G., Ronald Kahn, C. & Roth, J. Phlorizin: a review. Diabetes Metab. Res. Rev. 21, 31–38 (2005).

18. Duda, M. M., Varban, D. I., Muntean, S., Moldovan, C. & Olar, M. USE OF SPECIES AGASTACHE FOENICULUM (PURSH) KUNTZE. Hop Med. Plants 21, 52–54 (2014).

19. Soleimani, M., Arzani, A., Arzani, V. & Roberts, T. H. Phenolic compounds and antimicrobial properties of mint and thyme. J. Herb. Med. 36, 100604 (2022).

20. Shahdadi, F. et al. Mentha longifolia Essential Oil and Pulegone in Edible Coatings of Alginate and Chitosan: Effects on Pathogenic Bacteria in Lactic Cheese. Molecules 28, 4554 (2023).

21. Voigt, V., Franke, H. & Lachenmeier, D. W. Risk Assessment of Pulegone in Foods Based on Benchmark Dose–Response Modeling. Foods 13, 2906 (2024).

22. Faehnrich, B., Chizzola, R., Schabauer, A., Pracser, N. & Duerrschmid, K. Volatiles in dairy products after supplementation of essential oils in the diet of cows and influence on taste of cheese. Eur. Food Res. Technol. 243, 1783–1797 (2017).

23. Singh, D., Kumari, K. & Ahmed, S. Natural herbal products for cancer therapy. in Understanding Cancer 257–268 (Elsevier, 2022). doi:10.1016/B978-0-323-99883-3.00010-X.

24. Lorenzo, J. M. et al. Preservation of meat products with natural antioxidants from rosemary. IOP Conf. Ser. Earth Environ. Sci. 854, 012053 (2021).

25. Lin, Y., Shen, X., Yuan, Q. & Yan, Y. Microbial biosynthesis of the anticoagulant precursor 4-hydroxycoumarin. Nat. Commun. 4, 2603 (2013).

26. Bye, A. & King, H. K. The biosynthesis of 4-hydroxycoumarin and dicoumarol by Aspergillus fumigatus Fresenius. Biochem. J. 117, 237–245 (1970).

27. Liu, B., Raeth, T., Beuerle, T. & Beerhues, L. A novel 4-hydroxycoumarin biosynthetic pathway. Plant Mol. Biol. 72, 17–25 (2010).

28. Perez-Rodriguez, E., Aguilera, J. & Figueroa, F. L. Tissular localization of coumarins in the green alga Dasycladus vermicularis (Scopoli) Krasser: a photoprotective role? J. Exp. Bot. 54, 1093–1100 (2003).

29. Torres-Castillo, J. A. & Olazarán-Santibáñez, F. E. Insects as source of phenolic and antioxidant entomochemicals in the food industry. Front. Nutr. 10, 1133342 (2023).

30. Liu, J. et al. Effects of quercetin and daidzein on egg quality, lipid metabolism, and cecal short-chain fatty acids in layers. Front. Vet. Sci. 10, 1301542 (2023).

31. Houben, K. et al. Salicylic acid residues in products of animal origin. Food Risk Assess Eur. 2, (2024).

32. Xu, D. et al. The bacterial community and metabolome dynamics and their interactions modulate fermentation process of whole crop corn silage prepared with or without inoculants. Microb. Biotechnol. 14, 561–576 (2021).

33. Djoumbou Feunang, Y. et al. ClassyFire: automated chemical classification with a comprehensive, computable taxonomy. J. Cheminformatics 8, 61 (2016).

34. Feng, Z.-J. & Lai, W.-F. Chemical and Biological Properties of Biochanin A and Its Pharmaceutical Applications. Pharmaceutics 15, 1105 (2023).

35. Chen, Z., Ding, W., Yang, X., Lu, T. & Liu, Y. Isoliquiritigenin, a potential therapeutic agent for treatment of inflammation-associated diseases. J. Ethnopharmacol. 318, 117059 (2024).

36. Naz, S. et al. Chrysin: Pharmacological and therapeutic properties. Life Sci. 235, 116797 (2019).

37. Grochowski, D. M., Locatelli, M., Granica, S., Cacciagrano, F. & Tomczyk, M. A Review on the Dietary Flavonoid Tiliroside. Compr. Rev. Food Sci. Food Saf. 17, 1395–1421 (2018).

38. Petkowski, J. J., Seager, S. & Bains, W. Reasons why life on Earth rarely makes fluorine-containing compounds and their implications for the search for life beyond Earth. Sci. Rep. 14, 15575 (2024).

39. Curci, D., Sundaram, T. S., Ghidini, S. & Arioli, F. What We Know About per- and Polyfluoroalkyl Contamination Levels in Milk. A Review from the Last Decade. Foods 14, 2274 (2025).

40. Bonato, T., Pal, T., Benna, C. & Di Maria, F. Contamination of the terrestrial food chain by per- and polyfluoroalkyl substances (PFAS) and related human health risks: A systematic review. Sci. Total Environ. 961, 178337 (2025).

41. Food Detail - gouda cheese - PTFI. https://ptfi.markerlab.com/detail/food/GGB100362.

42. Food Detail - cow milk (vitamin d, pasteurized, homogenized) - PTFI. https://ptfi.markerlab.com/detail/food/GGB100035.

43. Periodic Table of Food Initiative. Metabolomics Analysis by SB-AQ LC/MS. (2024).

44. Knox, C. et al. DrugBank 6.0: the DrugBank Knowledgebase for 2024. Nucleic Acids Res. 52, D1265–D1275 (2024).

45. Groh, K., Geueke, B. & Muncke, J. FCCdb: Food Contact Chemicals database. Version 5.0. Zenodo 10.5281/ZENODO.4296944 (2020).

46. Kim, S. et al. PubChem 2025 update. Nucleic Acids Res. 53, D1516–D1525 (2025).

47. Rutz, A. et al. The LOTUS initiative for open knowledge management in natural products research. eLife 11, e70780 (2022).

48. Chandrasekhar, V. et al. COCONUT 2.0: a comprehensive overhaul and curation of the collection of open natural products database. Nucleic Acids Res. 53, D634–D643 (2025).

49. Gallo, K. et al. SuperNatural 3.0—a database of natural products and natural product-based derivatives. Nucleic Acids Res. 51, D654–D659 (2023).

50. The pandas development team. pandas-dev/pandas: Pandas. Zenodo 10.5281/ZENODO.13819579 (2024).

51. McKinney, W. Data Structures for Statistical Computing in Python. in 56–61 (Austin, Texas, 2010). doi:10.25080/Majora-92bf1922-00a.

52. Pedregosa, F. Scikit-learn: Machine Learning in Python. Mach. Learn. PYTHON.

53. Hunter, J. D. Matplotlib: A 2D Graphics Environment. Comput. Sci. Eng. 9, 90–95 (2007).

54. Waskom, M. seaborn: statistical data visualization. J. Open Source Softw. 6, 3021 (2021).

